# A complex mechanism translating variation of a simple genetic architecture into alternative life-histories

**DOI:** 10.1101/2024.01.05.574286

**Authors:** Jukka-Pekka Verta, Jacqueline E. Moustakas-Verho, Iikki Donner, Morgane Frapin, Annukka Ruokolainen, Paul V. Debes, Jaakko Erkinaro, Craig R. Primmer

## Abstract

Linking genes to traits is a central goal in biology. Despite progress in discovering genes associated with trait differences, a poor understanding of the functional mechanisms underlying genetic associations leaves us critically far from connecting genetic and phenotypic variation. This knowledge-gap is particularly large in multifaceted phenotypes of ecological relevance such as life-history traits. Using a multiomic dissection of the genotype-phenotype association in a large-effect maturation age gene - the transcription cofactor *vestigial-like 3* (*vgll3*) - in Atlantic salmon (*Salmo salar*), we show that *vgll3* mediates concerted changes of distinct molecular phenotypes associated with puberty in male gonads. *Vgll3* genotype conferring *early* maturity upregulates key genes controlling androgen production, cellular energy and adiposity, and TGF-β signaling, among others, thereby increasing the likelihood of earlier pubertal initiation. Genotype-dependent developmental trajectories are produced through VGLL3 interaction with distinct transcription factors, thus coordinating differential activation of regulatory pathways. These results reveal a genetically simple, yet functionally complex, architecture underlying alternative life-histories where variation in a single major effect gene produces pleiotropic variation in a spectrum of cellular traits. Our results further suggest that evolution in correlated phenotypes such as exemplified by alternative life history strategies may be mediated by a surprisingly simple genetic architecture.

## Introduction

A central challenge in biology is to understand how genetic differences alter fitness via function. Such endeavors often start by discovering the genetic basis of fitness-related phenotypes, yet in most cases the mechanisms connecting genotype to the phenotype remain unknown. Finding causal functional links between genetic variation and phenotypes is not only key for understanding the mechanisms that mediate evolutionary change, but it also unlocks phenotypes to be interpreted, treated, predicted, and bred.

Understanding of the genotype-phenotype map ^1^ for traits found in nature has been largely hindered by their complex genetic basis. Evolutionary theory predicts, however, that phenotypic evolution can be also mediated by single mutations of large effect ^2^. Indeed, large-effect genes have been found to underlie variation in adaptive traits of ecological relevance such as coloration ^3–6^, morphology ^7,8^, behavior ^9–11^, physiology ^12^, and life histories ^13–19^. However, what is largely missing, even for cases where single genes of large effect are known, is an understanding of how genotype variation influence phenotype variation via molecular function of the identified genes. This knowledge-gap is especially wide for functionally complex phenotypes such as life history traits where differences in physiology, development, and behavior tend to co-vary, where consequences of variation for fitness are large, and that have therefore often been expected to be highly polygenic ^20^.

For understanding the genotype-phenotype map in complex life histories, few species are as powerful as Atlantic salmon (*Salmo salar*). Salmon is among the most variable vertebrates on Earth in terms of life-history strategies ^21^, with a single major gene, the transcription cofactor *vestigial-like 3* (*vgll3*), encoding for nearly 40% of maturation age variation in natural populations ^14,22^. Genetic variation in *vgll3* associates with maturation age in humans as part of a more complex genetic architecture that hinders genotype-phenotype dissection ^23^. Genetic variation in *vgll3* is associated with an ensemble of puberty-related traits in Atlantic salmon including age at maturity ^14,22^, early male maturation ^24–26^, body condition ^25,27^, food acquisition preference ^28^, aerobic performance ^29^, aggressive behavior ^30^, and migration behavior ^31^. Although *vgll3* may represent the most pleiotropic large-effect life-history gene yet found, the molecular mechanisms by which *vgll3* genotypes have such pleiotropic effects on distinct phenotypes are not known.

We predicted that regulatory networks associated with *vgll3* mediate the genotype-phenotype associations and pleiotropy in this large-effect gene. Single ‘master regulators’ act as hubs in gene networks and can influence multiple phenotypes in a pleiotropic manner through their effects on co-regulated genes mediating development ^32,33^. We therefore hypothesized that the mechanism of genotype-phenotype associations in *vgll3* is likely linked to its function as a transcriptional regulator, as maturation age associates with the pre-pubertal expression of alternative isoforms of *vgll3* ^24^, in addition to nonsynonymous genetic differences in the gene itself ^14^. To test this, we studied the transcriptomic trajectory of the differentiating male gonad from early spring until breeding time in the autumn, and its association with *vgll3* genotype in a common-garden experiment. By using chromatin immunoprecipitation-sequencing to map VGLL3 gene regulatory elements, we further identify interacting transcription factors and show that VGLL3 plays a role in regulating key developmental processes in the gonad that are functionally connected to other traits beyond puberty, yet also associated with the *vgll3* genotype. The results shed light on the molecular machinery behind adaptive variation in maturation age and exemplify how single major-effect genes may alter multiple cellular phenotypes in a concerted manner to mediate the development of alternative life history strategies.

## Results

### Key pubertal mechanisms are upregulated in the *early* genotype of *vgll3*

Given the pronounced impact of *vgll3* genotype on maturation age, and its high expression in immature testes ^24^, we hypothesized that it serves as key regulator of maturation age prior to the onset of puberty. To test this, we assayed differences in gene expression (RNA-seq), gene regulatory elements, and VGLL3 binding regions (ChIPmentation) in immature testes of individuals with alternative homozygous *vgll3* genotypes conferring either late (*vgll3_LL_*) or early (*vgll3_EE_*) maturation, across the second year of growth of salmon juveniles (Fig 1A). We have previously shown that *vgll3* genotype association with maturation age was reproducible in controlled conditions and robust to population, ambient temperature, and husbandry conditions ^24–26^ (Supplementary analysis).

**Figure 1.**
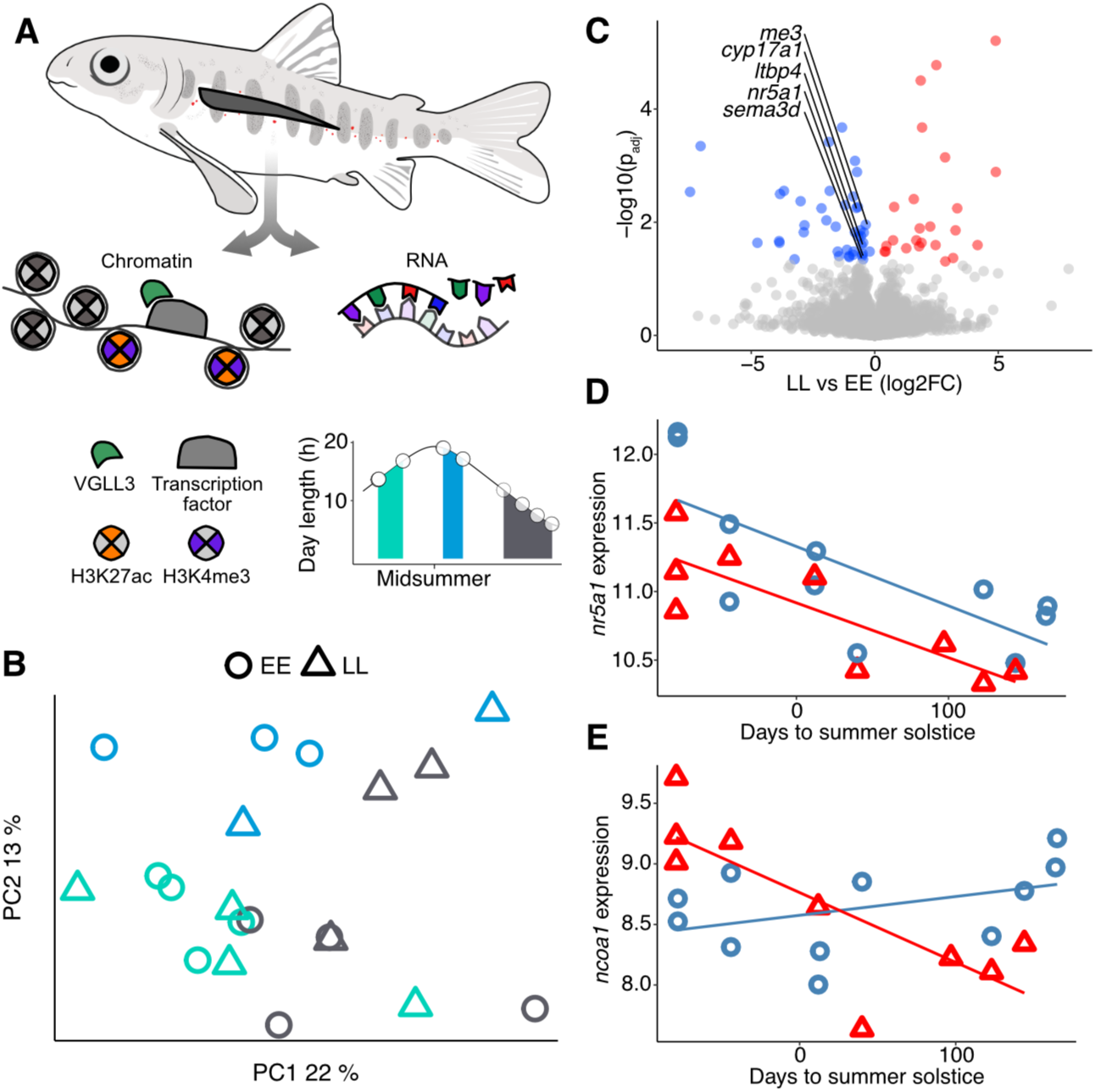
Identification of differentially expressed genes (DEGs) in immature salmon testes. A) Immature testes were sampled for RNA and chromatin across the second summer of growth. Sampling time points are indicated by white circles, with colored shading indicating the sampling period seasons. B) Testis transcriptomic data separate samples according to season. Samples are grouped in three categories for illustration purposes. C) DEGs between *vgll3* genotypes. Multiple key maturity genes show consistently higher expression in *vgll3_EE_* genotypes. D) *Nr5a1*shows higher expression in *vgll3_EE_* (blue) compared to *vgll3_LL_* (red) genotypes and both genotypes show a trend of decreasing expression towards autumn. E) *Ncoa1* expression shows an interaction between *vgll3* genotype (*vgll3_EE_;* blue, *vgll3_LL_*; red) and season.

Season was a major source of variation in testicular gene expression. Factor analysis ^34^ of testis RNA-seq data, which accounts for heterogeneity associated with e.g., cell cycle stage and sample maturation trajectories (Supplementary material), separated samples according to season based on the first two principal components of staged expression data (Fig 1B). Using a generalized linear model analysis, we identified 957 differentially expressed genes (DEGs) affected by sampling date, while adjusting for *vgll3* genotype (*p_ADJ_*<0.05). These DEGs included many TGF-β signaling and transcriptional regulator proteins, some of which show genetic association with pubertal age in humans ^23^ (Table S1). Our analyses therefore indicated that seasonal expression dynamics involve key signaling pathways in sexual development.

*Vgll3* genotype was associated with expression of a restricted, yet important, set of genes. We identified 70 DEGs between *vgll3* genotypes (*p_ADJ_*<0.05), while adjusting for season, with five genes of particular relevance to sexual maturation that were upregulated in *vgll3_EE_* genotype which confers early maturation, compared to typically later maturing *vgll3_LL_* individuals (Fig 1C); *nuclear receptor family 5 group A 1* (*nr5a1, a.k.a. SF-1,* Fig 1D), encoding for a transcription factor that controls the production of sex hormones and sexual development ^35^; *steroid 17-alpha-hydroxylase/17,20 lyase* (*cyp17a1*), a gene regulated by *nr5a1* that encodes for an essential step in androgen synthesis and is required for male maturation; *semaphorin 3D* (*sema3d*), encoding for a secreted signaling peptide that regulates cell adhesion and cytoskeleton during cell migration; *malic enzyme 3* (*me3*), where genetic and isozyme variation has been associated with growth ^36^ and maturity age variation in Atlantic salmon ^37^; and *latent-transforming growth factor beta-binding protein 4* (*ltbp4*), which regulates transforming growth factor beta (TGF-β) signaling. Concerted expression differences in genes associated with sex hormone production, cell migration, energy production, and TGF-β signaling pointed towards *vgll3* controlling for distinct facets of cellular puberty changes in the testes.

We additionally identified 35 DEGs with an interaction between *vgll3* genotype and sampling date (*p_ADJ_*<0.05), including *nuclear receptor coactivator-1* (*ncoa1* a.k.a. *steroid receptor coactivator-1* [*SRC-1*]). *Ncoa1* expression increased towards breeding time in *vgll3_EE_* individuals while it decreased in *vgll3_LL_* individuals (Fig 1E). Given that increasing *ncoa1* expression promotes lipid storage usage in mammals ^38^, our results suggest that, towards breeding time, the contrast in maturation age between *vgll3* genotypes translate into alternative dynamics of energy expenditure; *vgll3_EE_* individuals invest stored energy in processes enhancing puberty initiation through upregulation of *ncoa1* expression and subsequent lipid expenditure, while the relative lower expression of *ncoa1* in *vgll3_LL_* individuals predicts investment in adiposity and growth, and therefore future size and reproduction.

### Upregulation of maturation genes associates with VGLL3 binding

We reasoned that if functional differences in the VGLL3 protein mediate the observed expression differences, differentially expressed gene loci should be associated with VGLL3 binding. To test this, we performed ChIPmentation of histone modifications H3K27ac and H3K4me3, associated with active gene transcription, as well as VGLL3, based on a custom-produced antibody for Atlantic salmon.

We identified promoter regions as those associated with both H3K27ac and H3K4me3, while we defined enhancers as regions with H3K27ac alone ^39^; both features correlated with expression of nearest genes (Fig 2A-B). VGLL3 binding, when overlapping with promoters, associated with increased gene expression (Fig 2C, *p*<2.2e-16). *Vgll3* genotypes showed marked differences in both promoters and enhancers that were occupied by VGLL3 (Fig 2D-F). VGLL3 binding was observed in a core set of promoters and enhancers in both genotypes (VGLL3 promoters: observed 2747, expected 97, *p*=0.009, VGLL3 enhancers: observed 11137, expected 1172, *p*=0.009), with nearly equal numbers of regulatory elements with VGLL3 occupancy specific to each genotype (Fig 2E). To test for functional effects of this genotype difference in VGLL3 binding, we analyzed GO overrepresentation in each category relative to all VGLL3 regulatory elements (promoters and enhancers). Genes associated with VGLL3 elements unique to the *vgll3_EE_* genotype were overrepresented in signaling receptors, cell adhesion genes, and those involved in actin regulation, among other functions (*FDR*>0.1, Fig 2F). Genes associated with VGLL3 elements unique to *vgll3_LL_* were overrepresented in semaphoring receptors and regulators of cell cycle progression, among others (*FDR*>0.1, Fig 2F). Interestingly, cell differentiation process was over-represented independently in both genotypes, indicating that VGLL3 regulatory loci influencing this process were largely unique to each genotype, or that they were acting as promoters in one genotype and enhancers in the other. Taken together, these results indicate that VGLL3 is widely associated with gene regulatory regions in immature testes and suggest that *vgll3* genotype differences have wide-scale influence on cellular physiology and development through coordinated regulation of distinct genomic loci and cellular functions.

**Figure 2.**
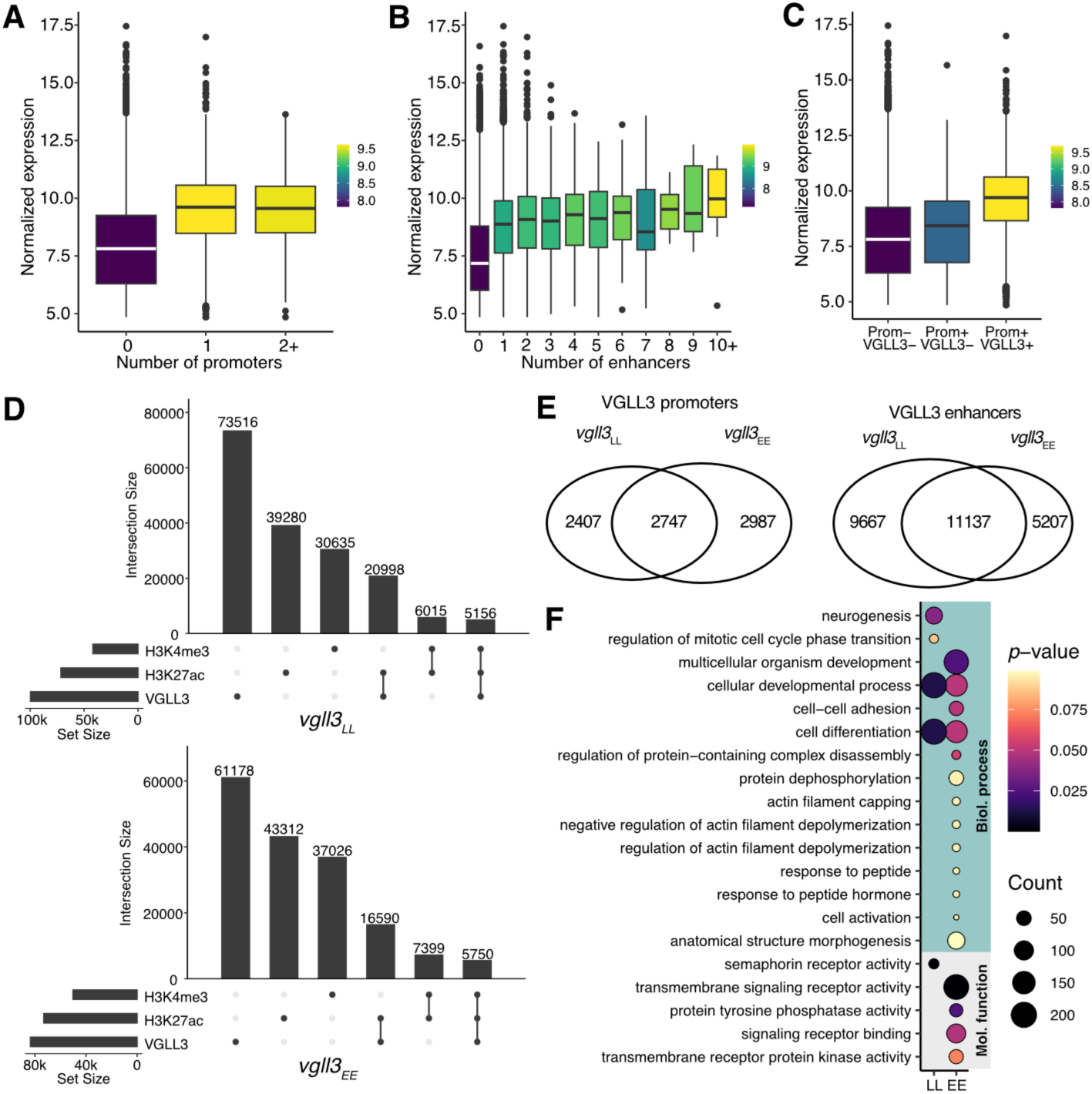
Identification of gene regulatory elements by genome-wide mapping of histone modifications and VGLL3 binding regions. The number of A) promoters (H3K27ac + H3K4me3) and B) enhancers (H3K27ac alone), assigned to nearest expressed genes correlate positively with gene expression levels. C) VGLL3 binding, when associated with promoters, is associated with increased gene expression. D) Numbers of VGLL3 regions, promoters, and enhancers in *vgll3* genotypes. E) Sharing of VGLL3 promoters and VGLL3 enhancers between *vgll3* genotypes (log_FC_>2). F) Over-represented gene ontologies in genes with VGLL3 promoters or VGLL3 enhancers that are unique to each genotype (adjusted *p-*value > 0.1).

We proceeded with identification of regulatory elements that could directly mediate the *vgll3* genotype effect on gene expression and further on maturation age. In total, 50 of the 70 DEGs between *vgll3* genotypes were associated with VGLL3 binding regions (23 with VGLL3 regions within promoters or enhancers), suggesting that expression differences may be directly mediated by functional differences in VGLL3 protein. DEGs with VGLL3 promoters or enhancers showed a notable trend with the majority having higher expression in *vgll3_EE_* individuals (5/5 VGLL3 promoters, 17/23 VGLL3 enhancers), suggesting that transcriptional upregulation associated with the VGLL3_EE_ protein was stronger compared to the VGLL3_LL_ protein. The upregulated DEGs included *nr5a1, cyp17a1, me3, sema3d,* and *ltbp4,* each associated with a central process in sexual maturity, namely, androgen production, metabolism, cell motility, and TGF-β signaling (Fig 3A-B). Of these, VGLL3 binding in proximity of *nr5a1, cyp17a1,* and *sema3d* associated with differential H3K4me3 enrichment, with histone trimethylation observed only in *vgll3_EE,_* but this signal was driven by strong H3K4me3 enrichment in a single sample. Overall, our results strongly suggested that VGLL3 binding mediated the higher expression of key maturation genes in the *vgll3_EE_* genotype, corresponding to a concerted upregulation of signaling pathways pre-puberty.

**Figure 3.**
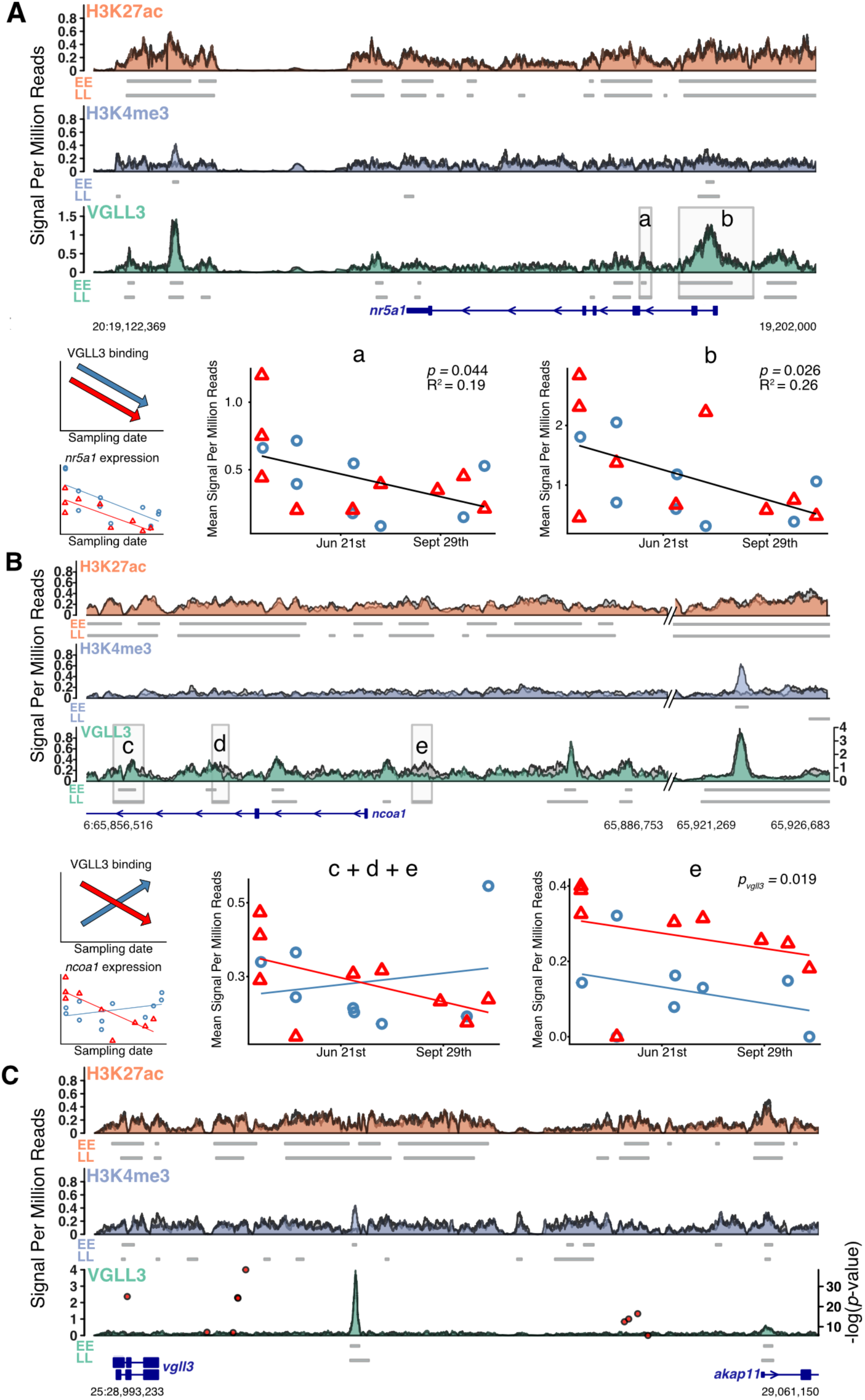
VGLL3 binding is associated with *vgll3* genotype effects and seasonal expression variation in central maturation genes. ChIPmentation signals from *vgll3_EE_* and *vgll3_LL_* genotypes are overlaid for each histone modification/VGLL3 (*vgll3_LL_* on grey). Horizontal bars below ChIPmentation signal represent enriched regions called by MACS3 in each genotype. (A) Decreasing *nr5a1* expression towards the autumn is putatively mediated by two VGLL3 regulatory regions (shaded areas; a & b) in proximity of the *nr5a1* promoter that show a decrease of VGLL3 binding signal towards the autumn. (B) Opposing *ncoa1* expression patterns of *vgll3* genotypes across the season are putatively mediated by VGLL3 binding in three regulatory regions in proximity of the *ncoa1* promoter (shaded areas; c, d & e). VGLL3 binding in region *e* shows strong differences between *vgll3* genotypes. (C) Intergenic region adjacent to the *vgll3* locus shows a prominent VGLL3 binding region. GWAS association *p-*value from ref (*15*) is represented as red points on VGLL3 ChIP-signal background (axis on left).

To test if VGLL3 binding could mediate some of the most salient patterns of seasonal expression variation, namely the decrease in expression of *nr5a1* towards the autumn, we investigated the strength of VGLL3 binding signal in the proximity of the *nr5a1* promoter. We identified two VGLL3 binding regions that showed a decrease in VGLL3 signal towards the autumn, indicating that they likely mediated *nr5a1* expression dynamics (Fig 3A). We further identified proximal VGLL3 binding regions (7 VGLL3 enhancers) in 16 DEGs showing interaction between *vgll3* genotype and season. Our expression data showed that seasonal expression of *ncoa1* was dependent on *vgll3* genotype; a declining expression in *vgll3_LL_* genotype suggested investment on adiposity, while an increasing trend in *vgll3_EE_* genotypes was suggestive of utilization of fat reserves to promote pubertal initiation. VGLL3 signal in one VGLL3 enhancer in proximity of *ncoa1* showed strong differences between *vgll3* genotypes; when signal was averaged over two additional VGLL3 enhancers a tentative pattern of interaction between season and *vgll3* genotype was observed, albeit the trend was statistically non-significant (Fig 3B). An additional striking VGLL3 binding region having one of the strongest signals in the genome was observed in the first intron of *ncoa1,* and associated with differential H3K4me3 in *vgll3* genotype, albeit the H3K4me3 signal that was driven by a single sample. Overall, our results identify specific VGLL3 regulatory elements that likely mediate expression differences of key downstream genes between *vgll3* genotypes, season, and their interaction.

### *vgll3* self-regulation as a genotype-phenotype link in maturation age

In natural populations, the strongest association between maturation age and genetic variation was observed in a non-coding region adjacent to *vgll3* ^14^, suggesting that divergence in gene regulatory elements acting in *cis* to *vgll3* contribute to the causal mechanism of maturation age variation. To test this, we used testis chromatin modification and VGLL3 binding data, and identified a prominent VGLL3 binding region adjacent to the analyzed SNP of strongest association signal (Fig 3D) ^14^. VGLL3 binding was observed for both genotypes but an overlapping H3K4me3 region was observed in *vgll3_EE_* but not in *vgll3_LL_* individuals; this pattern was driven by strong histone trimethylation in a single sample. Taken together, these results suggest that the genotype-phenotype association between *vgll3* and maturation age includes differential self-regulation of the *vgll3* gene by itself, possibly forming a feedback loop that might interplay with the non-synonymous mutations and splicing variation also associated with maturation age ^14,24^. Overall, our results support a mechanism where strong *trans* effects on gene networks are underlined by *cis* variation in regulatory genes themselves ^32^.

### VGLL3 drives gene co-expression differences in diverse signaling pathways

Single master regulators may drive differences in complex phenotypes by way of influencing the expression of a co-regulated set of genes ^32^. To test if *vgll3* could mediate its effects on molecular phenotypes in this way, we investigated the effect of *vgll3* genotype and season on gene regulatory networks using gene co-expression network analysis. Expressed genes were assigned to 70 co-expression modules ranging in size from 31 to 10972 genes. Using an ANOVA model to test the difference in module (eigengene) expression between *vgll3_EE_* and *vgll3_LL_* genotypes, we identified four co-expression modules with higher expression in *vgll3_EE_* individuals (*P*<0.05, not significant after correction for multiple testing, Fig 4A). Module genes were connected to DEGs by function and involved in cell adhesion, immune system, chemokine and cytokine signaling, TGF-β signaling, and Hippo signaling (Fig S3). Notably, the *magenta* module supported a regulatory link between Hippo and Ras signaling mediated by VGLL3 binding (Fig 4B, Supplemental analysis). Additionally, one module showed higher expression in *vgll3_LL_* individuals; it was over-represented in genes associated with cell-cell signaling and negative regulation of the Wnt pathway (e.g., *APC regulator of WNT signaling pathway*), further supporting that *vgll3* genotype mediated concerted changes in cell adhesion and motility through gene co-regulation.

**Figure 4.**
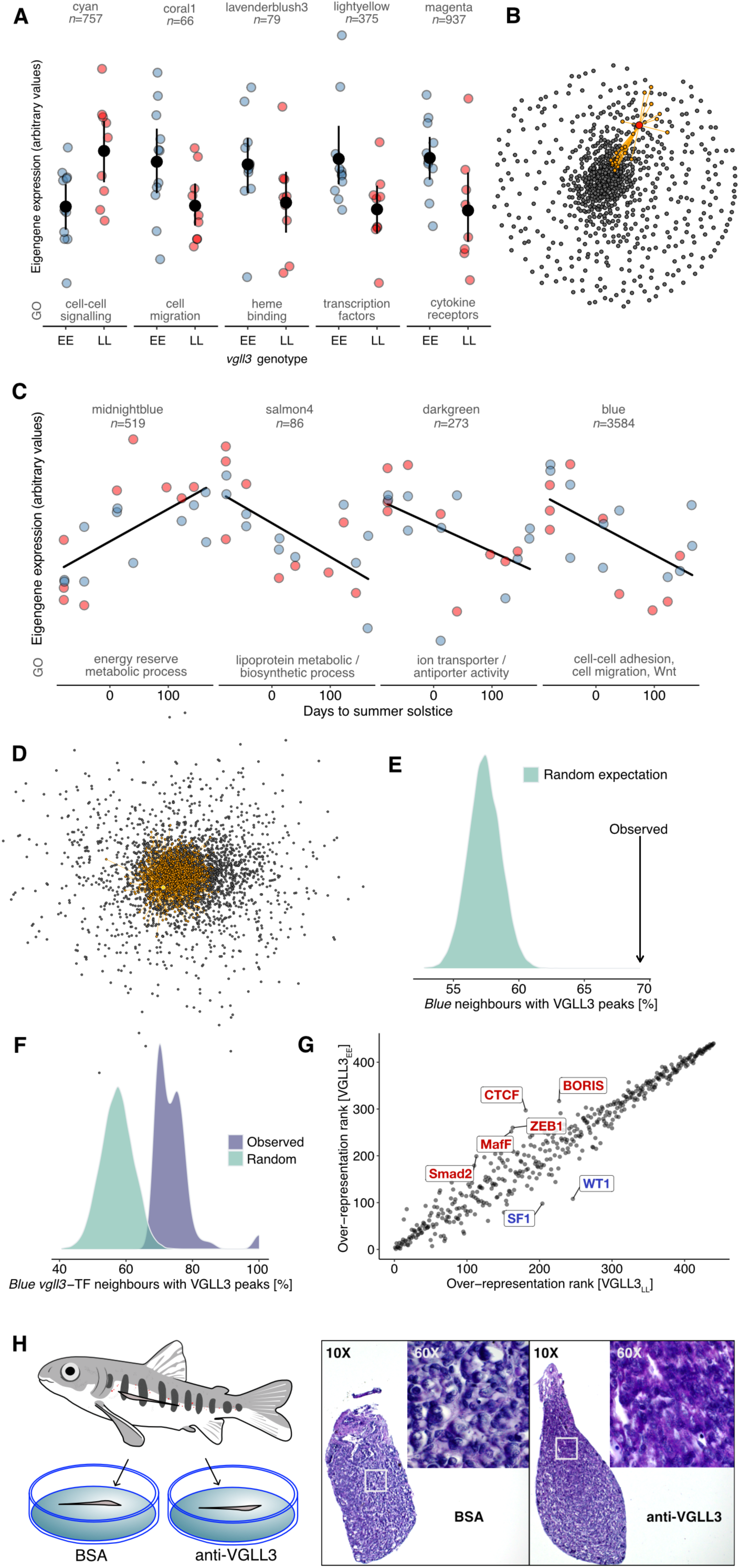
Gene co-expression networks reveal signaling pathways associated with *vgll3* genotype and seasonal variation in testes transcriptomes. A) Five modules show significant eigengene expression difference between *vgll3* genotypes (*p*<0.05, not significant after correction for multiple testing). Modules are annotated according to the number of included genes and a selected, overrepresented GO category. Blue spheres, *vgll3_EE_* individuals; red spheres, *vgll3_LL_* individuals; black spheres, mean; whiskers, 95% confidence limit estimates B) The *magenta* module includes the known *vgll3* molecular partner *tead3* (red sphere) and its neighbors (TOM>0.2, orange spheres) include the Ras signaling inhibitor *neurofibromin*. Genes showing higher correlation in expression are situated towards the center of the network. C) Expression of four modules show significant association with season (*p_ADJ_*<0.05). D) The *blue* module includes *vgll3* (golden sphere). *Vgll3* neighbors (TOM<0.2) are represented as orange spheres. Genes are ordered according to the strength of their co-expression, showing that *vgll3* expression is highly correlated with most of the genes in *blue* (*vgll3* situates towards the center of the network). E) Percentage of *blue* module neighbors with proximal VGLL3 binding regions (arrow), compared to expected distribution based on random sampling of genes. F) Percentage of shared neighbors between *vgll3* and transcription factors included in *blue* with proximal VGLL3 binding regions, compared to expectation from random sampling of genes. G) Difference in TF motif over-representation in *vgll3_LL_* and *vgll3_EE_* enhancers and promoters, compared to all enhancer and promoter regions within genotypes. Over-represented motifs are rank-transformed from most highly over-represented to least over-represented. Motifs with a rank difference over 80 are highlighted. H) Tissue sections of immature testis from the same individual show that knock-down of VGLL3 function induces cellular proliferation and motility. Top right corners show 60X magnification of area outlined with white rectangles.

Four co-expression modules showed a significant correlation with sampling date (*p_ADJ_*<0.05, Fig 4C). Among the three modules with decreasing trend towards spawning time, the *blue* module showed strong over-representation in genes involved in cell adhesion and morphogenesis, cell motility, and Wnt signaling, among others (Fig S4). *Blue* included *vgll3*, *vgll4*, *tead1b*, *tead2*, *nr5a1*, *cyp17a1*, *me3*, *sema3d*, *Wnt4*, among other genes (Table S3). *Blue* module expression pattern was consistent with a decreasing expression of genes inhibiting epithelial-to-mesenchymal transition (EMT, e.g., *cadherin*, *protocadherin*, and *Wnt4*) towards the breeding season, consistent with a role of *vgll3* in the regulation of EMT ^40^. *Vgll3* was among the most connected genes within *blue* with 1490 neighbors of the 3563 genes in *blue* in total (75^th^ quantile, Fig 2D). *Blue* also contained 103 transcription factor genes (GO:0003700), which allowed us to identify putative regulatory partners of *vgll3* (Fig S5). Top transcription factors with most network neighborhood sharing with *vgll3* included immune system, retinoic acid, Hippo, homeobox, thyroid hormone, and nuclear factor regulators, putatively connecting *vgll3* to diverse cellular signaling pathways including membrane-bound receptors and nuclear receptors (Fig S5); these gene pathways have been implicated in maturation age variation in humans as well^23^.

We next investigated VGLL3 binding in proximity of *vgll3* network neighbors in the *blue* module to test if VGLL3 binding could mediate the co-regulation of the genes. Of all *vgll3* neighbors, 69% were associated with VGLL3 binding regions, (significantly more than expected by chance, Fig 4E), suggesting that *blue* co-expression could largely be mediated by regulation by VGLL3. Similarly, shared network neighbors between *vgll3* and transcription factors in *blue* were overrepresented in genes with VGLL3 binding regions compared to a random expectation (Fig 4F), suggesting that *blue* co-expression was mediated by VGLL3 interaction with a diverse set of transcription factors.

### *vgll3* genotypes associate with distinct transcription factor co-occupancies

Our results indicated that *vgll3* genotypes were associated with large-scale differences in gene expression and regulatory element activity, and that these differences were mediated by VGLL3 binding. VGLL3 does not contain a DNA-binding domain, rather, it interacts with transcription factors to regulate gene expression ^41^. We used motif analysis of VGLL3 binding regions to identify transcription factors that putatively co-occupy regulatory elements with VGLL3 and may thus mediate the genotype effect on gene regulation. Although motif over-representation was largely conserved in VGLL3 regions between the genotypes, notable differences in motif enrichment indicated that VGLL3 co-occupied regulatory elements with distinct sets of transcription factors in different genotypes (Fig 4G). Motifs with most pronounced over-representation in *vgll3_EE_,* compared to *vgll3_LL_*, were *nr5a1* (*SF-1*) and *WT1*, both positive regulators of sexual development and maturation. In *vgll3_LL_*, compared to *vgll3_EE_*, most over-represented motifs corresponded to *ZEB1, OVOL2, SMAD2, BORIS, CTCF,* and *MafF*, involved in the regulation of epithelial-mesenchymal transition (EMT), inhibition of androgen receptor activity, and chromatin structure. These results indicate that VGLL3 interacts with distinct sets of transcription factors that vary depending on the *vgll3* genotype; either promoting sexual maturation in *vgll3_EE_*, or putatively inhibiting it in *vgll3_LL_*.

### Knock-down of *vgll3* induces pubertal-like cell differentiation

Our genomic data suggested that *vgll3* controls cell differentiation and motility through aligned regulation of multiple cellular signaling cascades, especially *nr5a1-ncoa1,* Hippo-Ras, and TGF-β signaling. To test the causative role of *vgll3* in mediating cell proliferation, differentiation, and motility changes, we cultured immature testes in the presence of anti-VGLL3 antibody, thus knocking down *vgll3* gene function. Knock-down of VGLL3 led to notable changes in testis morphology and cellular phenotypes; compared to control testis from the same individuals, knock-down testis exhibited increased cell density (likely from cell proliferation), as well as increased cell motility that were evident as protruding masses of cells (Fig 4H, Fig S6, *n*=8). These results provide strong support for the hypothesis that *vgll3* acts as a central regulator controlling the initiation of maturation-associated cellular changes in the testis through distinct yet connected cellular processes.

## Discussion

A dissection of the proximal mechanisms linking genes to traits is required for a complete understanding of evolutionary change ^1^. Despite the simple genetic architecture of maturation age variation in Atlantic salmon, we show here that the mechanism acting through the transcription cofactor *vgll3* integrates the regulation of multiple distinct signaling pathways and developmental programs, thus being functionally multifaceted.

Our results show that the association between *vgll3* genotype and maturation age in Atlantic salmon represents a case of a genetic effect with a mechanism we propose as “sub-complex”. In contrast to a simple mechanism such as documented in e.g., coloration ^3–6^, salmon maturation age seems to have a simple genetic basis that through regulatory pleiotropy influences multiple traits. It can thus be thought of as a single “sub-unit” of a more complex genetic architecture (Fig 5). We anticipate that additional genes mediating the diverse cellular functions regulated by *vgll3* explain the remaining polygenic component of maturation age acting in addition to the gene itself ^25,42^. The complexity of mechanisms discovered here also imply that in species where life history variation has a more complex genetic architecture, such as maturation age variation has in humans ^23^, we can expect an extremely complex genotype-phenotype map approaching to what has been proposed for other complex traits ^32^. Overall, our results exemplify a hidden complexity of the molecular mechanisms mediating the large, pleiotropic effect of a single gene on alternative life histories.

**Figure 5.**
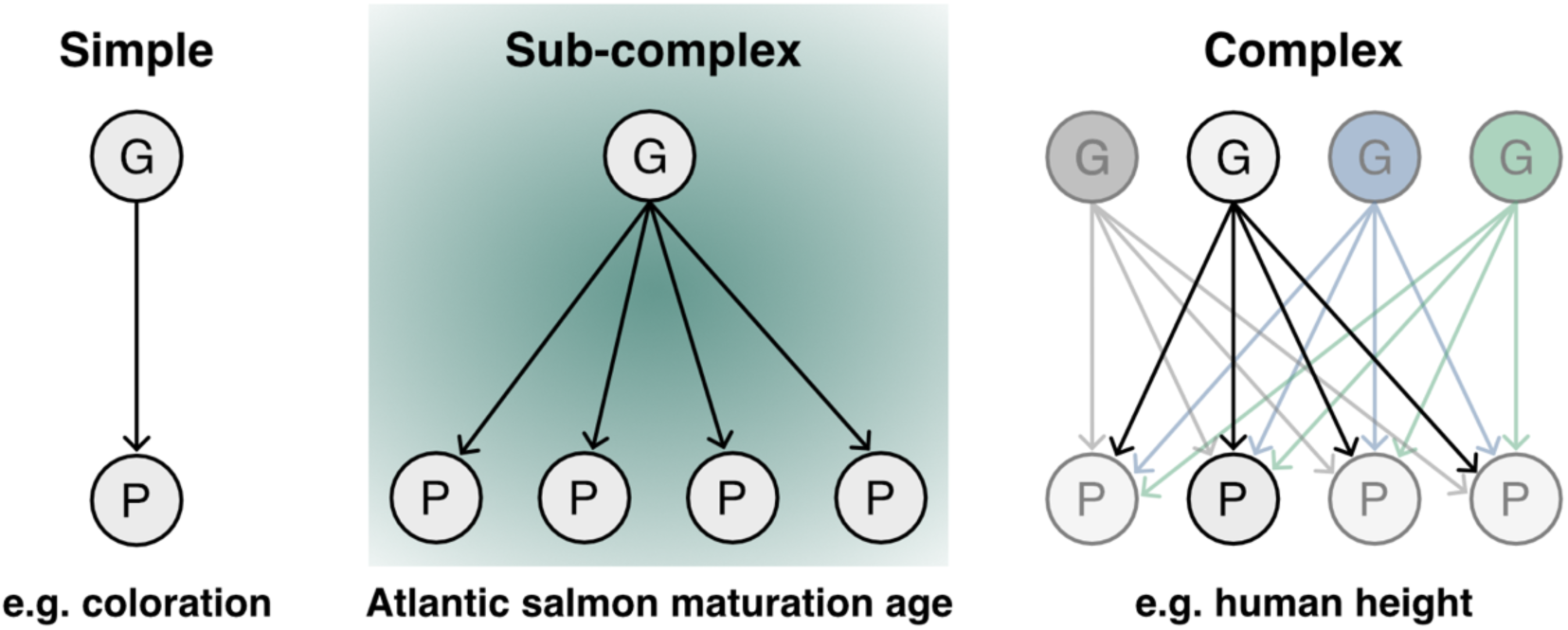
A proposed model for the architecture of the genotype (G) to phenotype (P) map in different traits. Single genes with pleiotropic effects on multiple phenotypes, such as *vgll3* in Atlantic salmon, are proposed to represent single “sub-units” of complex genetic architectures where single genes have smaller effects.

Our results also bring significant insight into the molecular basis of pleiotropy in large effect genes by showing that *vgll3* is connected to diverse physiological and behavioral phenotypes through its central role in gene regulation. We propose that by modulating the interaction with different transcription factors, *vgll3* genotypes mediate correlated changes in the regulation of distinct downstream signaling pathways. For example, changes in *nr5a1*/*nr5a2* signaling can cause absence of puberty or variation in maturation age in humans ^43^ and in rainbow trout (*Oncorhynchus mykiss*) ^18^; *nr5a1* also regulates aggressivity and physical activity ^44–46^, traits that are linked to genetic variation in *vgll3* in salmon ^30,31^. Body condition is another trait linked to *vgll3* in both salmon and humans ^25,27,47^, as is metabolic rate and aerobic scope ^29^. Here, the likely functional link acts through *vgll3* regulation of *ncoa1* expression, as ablation of *ncoa1* expression lowers cellular oxygen consumption and leads to body weight gain in mice ^48^, corroborating with seasonal changes in salmon body condition that depend on *vgll3* genotype ^27^. Our results therefore showcase that changes in regulatory interactions of a single gene can mediate phenotypic covariation in a spectrum of seemingly unrelated traits. In the light of this, evolution in functionally complex phenotypes such as alternative life-histories and the “pace-of-life”, typically considered to be highly polygenic characteristics of species, may indeed be mediated by a surprisingly simple genetic architecture.

## Acknowledgements

We thank Ana Lindeza, Claudius Kratochwil, and Frédéric Guillaume for comments on the manuscript, Nikolai Piavchenko and Noora Parre for fish husbandry and Seija Tillanen for help with lab work.

## Funding

Funding was provided by Academy of Finland (grant numbers 314254, 314255, 327255 and 342851), the University of Helsinki, and the European Research Council under the European Articles Union’s Horizon 2020 and Horizon Europe research and innovation programs (grant no. 742312 and 101054307). Views and opinions expressed are however those of the author(s) only and do not necessarily reflect those of the European Union or the European Research Council Executive Agency. Neither the European Union nor the granting authority can be held responsible for them.

## Author contributions

Conceptualization J.-P.V., C.R.P., P.V.D., Formal analysis J.-P.V., Methodology J.-P.V., J.M.-V., P.V.D., Investigation J.-P.V., J.M.-V., I.D., M.F., A.R., Resources C.R.P., J.E., Supervision J.-P.V., C.R.P., Writing – original draft J.-P.V., Writing – review & editing J.-P.V., C.R.P.

## Data availability

Sequence data is publicly available in NCBI Sequence Read Archive upon publication (PRJNA1042649). Output files for gene expression and chromatin features as well as all R code used in analysis is publicly available in Dryad (https://doi.org/10.5061/dryad.vhhmgqp1g).

## Material and Methods

### Material

Fish crosses used are described in depth in ref ^24^. Briefly, 2×2 factorial matings between *vgll3* LL and EE individuals were used to create 32 families of *vgll3* LL, LE, EL and EE individuals. The broodstock is a first-generation hatchery stock composed of a mixture of northern Baltic Sea lineages and maintained by the Natural Resources Institute Finland (LUKE). Eggs were fertilized in October 2017 and raised in controlled conditions at the University of Helsinki research animal facility. Two egg incubators with families separated by mesh compartments were used for housing eggs and alevin up to first feeding. After first feeding, 48 individuals were randomly selected from each family and distributed in roughly equal densities across four 0.25 m^3^ recirculating tanks.

Fish were raised using a typical annual cycle of water temperature and photoperiod corresponding to latitude of origin (min-max temp: 6.3-17.7 °C, latitude: 61N) for 1,5 years with *ad libidum* feeding of appropriately sized commercial fish feed. At eight months of age, fish were individually tagged with passive integrated transponders (PIT-tags) and a fin clip was collected for genotyping. Genotyping was performed using a set of 141 SNP markers, including *vgll3*, and PCR-sequencing ^49^. Individuals were assigned to families using SNPPIT ^50^.

Individuals were pre-assigned randomly for sampling dates with equal representation of genotypes and sexes. At assigned time points, fish were euthanized using an overdose of buffered tricaine methanesulphonate (MS-222) and dissected. Their maturity phenotype was recorded and tissue samples including testes were flash-frozen on liquid nitrogen and stored in −80 °C until extraction of RNA and chromatin. Testis tissue for ChIPmentation was immersed in 1 ml Cryostor10 in cryovials and incubated on ice. Cryovials were transferred to a MrFrosty freezing container and transferred to −80 °C. Cryostored tissue was stored in −80 °C until extraction of cells for chromatin. Experimentation was conducted according to the Finnish Government Decree on the Protection of Animals Used for Scientific or Educational Purposes (564/2013), which implements EU directive 2010/63/EU. The experiments in this study were approved by the Project Authorisation Board (ELLA) on behalf of the Regional Administrative Agency for Southern Finland (ESAVI) under experimental license ESAVI/2778/2018.

### RNA extraction and RNA-seq

Gonad tissue was removed from −80 °C storage and flash-frozen in Macherey-Nagel (MN) Nucleospin 96 RNA extraction buffer. Frozen samples for 9 *vgll3_LL_* individuals and 10 *vgll3_EE_* individuals were homogenized on a OMNI Bead Ruptor Elite instrument using 2 ml tubes with 2.4 mm steel beads and a program with 6 bursts of 20s with speed 4,5 and 10s pauses in between bursts. Total RNA was extracted following MN kit instructions on 96-well plates. RNA elutions were treated with Invitrogen Turbo DNase kit reagents for removal of trace DNA and quantified using Thermo Quant-iT reagents on a 96-well plate reader. RNA concentration was verified using a Qubit instrument and Quant-iT reagents, and adjusted to 50 ng/μl with pure water. 100 ng of total RNA was used for RNA-seq library construction using Illumina stranded mRNA kit and manufacturer instructions. Libraries were sequenced at the University of Helsinki Institute of Biotechnology sequencing service using a NextSeq500 instrument and 75 bp paired-end reads.

### Antibodies

A custom anti-VGLL3 polyclonal antibody was procured from Genscript with delivery in January 2020 as follows. Target DNA sequence of *vgII3* was optimized and synthesized. The synthesized sequence was cloned into vector pET-30a (+) with His tag for protein expression in *E. coli*. *E. coli* strain BL21 star (DE3) was transformed with recombinant plasmid. A single colony was inoculated into TB medium containing related antibiotic; culture was incubated in 37 °C at 200 rpm and then induced with IPTG. SDS-PAGE was used to monitor the expression. Recombinant BL21 star (DE3) stored in glycerol was inoculated into TB medium containing related antibiotic and cultured at 37 °C. When the OD600 reached about 1.2, cell culture was induced with IPTG at 37 °C/4h. Cells were harvested by centrifugation. Cell pellets were resuspended with lysis buffer followed by sonication. The precipitate after centrifugation was dissolved using denaturing agent. Target protein was obtained by one-step purification using Ni column. Target protein was sterilized by 0.22 μm filter before stored in aliquots. The concentration was determined by BCATM protein assay with BSA as standard. The protein purity and molecular weight were determined by standard SDS-PAGE along with Western blot confirmation. Two rabbits were immunized with three injections of purified VGLL3 protein and antiserum was tested for presence of anti-VGLL3 antibodies after 3^rd^ bleeding using Western blotting of Atlantic salmon heart tissue. Rabbits were immunized for a fourth time, sacrificed, antiserum was pooled, and anti-VGLL3 antibody was affinity-purified from the antiserum pool. Purified VGLL3 antibody was validated for reactivity using an indirect ELISA assay VGLL3 protein as antigen.

Additional quality-control for the VGLL3 antibody was performed using co-immunoprecipitation followed by Western blotting of Atlantic salmon heart tissue (Fig S7), which was shown to have high *vgll3* expression ^24^, as follows. The VGLL3 antibody was coupled with beads from the Dynabeads Co-Immunoprecipitation Kit (Invitrogen, Massachusetts, USA) using 10 μg antibody per mg beads. Protease inhibitors were added to the Extraction Buffer and flash-frozen salmon hearts weighing approximately 0,25 g were homogenized in 2250 μl buffer each using the Bead Ruptor Elite (Omni International, Georgia, USA). The lysates were centrifuged at 840 x g for two minutes in 4 °C and the supernatants combined with 150 μl antibody-coupled beads. Proteins and beads were then incubated at 4 °C for 40 minutes. The rest of the co-immunoprecipitation was performed in accordance with manufacturer protocol.

The co-immunoprecipitated protein samples and positive controls (purified VGLL3 protein) were prepared for western blotting by adding appropriate amounts of 4X laemmli and DTT (final concentration of DTT 100 mM) and incubating at 92 °C for five minutes. Proteins were separated on Any kD Mini-PROTEAN TGX Precast Protein Gels in a mini electrophoresis system (Bio-Rad Laboratories, California, USA). For molecular weight sizing, Precision Plus Protein WesternC Standards (Bio-Rad Laboratories, California, USA) were used. Separated proteins were transferred to a 0,2 µm nitrocellulose membrane using Trans-Blot Turbo Transfer Packs and the Trans-Blot Turbo Transfer System (Bio-Rad Laboratories, California, USA). 3% BSA in Tris-buffered saline with 0,1% Tween 20 (TBST) was used for membrane blocking and for diluting the primary antibody 3:1000. The incubation of the membrane in primary antibody was performed O/N at 4 °C. For protein detection, we used mouse anti-rabbit IgG-HRP sc-2357 (Santa Cruz Biotechnology, Texas, USA) and Precision Protein StrepTactin-HRP Conjugate (for protein standards; Bio-Rad Laboratories, California, USA), both diluted 1:20 000 in TBST. Chemiluminescence was recorded with the Pierce ECL Western Blotting Substrates (Thermo Fisher Scientific, Massachusetts, USA) and the ChemiDoc MP imaging system (Bio-Rad Laboratories, California, USA).

### Chromatin extraction and ChIPmentation

Detailed ChIPmentation protocol is provided as a supplement. In summary, gonad tissue for individuals used in RNA-seq was removed from −80 °C storage and rapidly thawed in a 37 °C water bath. Thawed tissue was rinsed with ice-cold D-PBS and transferred to a clean tube. Tissue was homogenized in ice-cold D-PBS using a OMNI Bead Ruptor Elite instrument with 7 ml tubes and 2.4 mm ceramic beads, and a program with one burst of 5s with speed 2.4. Cell suspension was filtered through a Flowmi Cell Strainer and inspected under a microscope with Trypan blue staining. Cell suspension was diluted with room temperature D-PBS and cross linked with Diagenode ChIP crosslinking gold in 1X concentration for 30 min, followed by fixation with 1% formaldehyde for 2 min. Formaldehyde was quenched in 0.125 M glycine for 5 min and cells collected with centrifugation at 400 g for 10 min. Cells were washed two times with 500 μl ice-cold PBS and centrifuged at 400 g for 10 min in between washes. Cells were subject to ChIPmentation with Thermo MAGnify ChIP-kit and Illumina Tn5 reagents as detailed in the supplement and briefly below.

Cells were collected using centrifugation, resuspended in 50 μl lysis buffer supplemented with protease inhibitors and lysed on ice for 5 min. Chromatin was sheared in 50 μl volumes using a Bioruptor device with settings high power and 3x eight cycles of 30s on, 30s off. Debris was pelleted by centrifugation and sheared chromatin was diluted to four equal aliquots of 100 μl using dilution buffer supplemented with protease inhibitors. One aliquot of sheared chromatin was reserved as input control. Remaining three aliquots were immunoprecipitated in 4 °C o/n using 1μg of Abcam ab4729, 2μg of Abcam ab8580, and 10 μg of a custom procured anti-VGLL3 antibody on ThermoFisher Dynabeads Protein A/G. Beads were subsequently washed following MAGnify kit protocol, with an additional final wash using 150 μl of ice-cold 10mM Tris (pH 8). Bead-bound chromatin was then treated in 20 μl volume of tagmentation reaction containing Illumina Tn5 transposase for 5min at 37 °C. Input controls were treated with tagmentation reaction for 5min at 55 °C. Tagmentation was terminated by adding 7.5 volumes of RIPA buffer and incubation on ice for 5min. Chromatin was subsequently washed twice with 150 μl of ice-cold RIPA and TE buffer. Crosslinks were reversed using a proteinase-K treatment and ChIPmentation DNA was captured using Macherey-Nagel NucleoMag magnetic beads. ChIPmentation libraries were measured using a Qubit instrument and a control-PCR was run with Nextera sequencing oligos to assess library amplification on agarose gel. Finally, libraries were indexed, pooled, and sequenced at the University of Helsinki Institute of Biotechnology sequencing and FIMM sequencing services using NextSeq500 (75 bp paired-end) and Novaseq6000 (150 bp paired-end) instruments, respectively.

### RNA-seq analysis

RNA-seq reads were trimmed using *fastp* ^51^ version 0.20.1 and options *p trim_front1=2 trim_front2=2 detect_adapter_for_pe*. Trimmed reads were aligned using *STAR* ^52^ version 2.7.9a and manual two-pass mode to the Atlantic salmon genome (Salmo_salar-GCA_905237065.2) downloaded from Ensembl. Alignment options for *STAR* were defined as *outFilterIntronMotifs RemoveNoncanonicalUnannotated chimSegmentMin 10 outFilterType BySJout alignSJDBoverhangMin 1 alignIntronMin 10 alignIntronMax 1000000 alignMatesGapMax 1000000 alignEndsProtrude 10 ConcordantPair limitOutSJcollapsed 5000000*. Reads overlapping Ensembl gene models (Salmo_salar-GCA_905237065.2) were quantified using R (version 4.2) package *Rsubread* ^53^ and the function *featureCounts*, specifying the parameters *countChimericFragments=FALSE, countReadPairs=TRUE, countMultiMappingReads=TRUE, fraction=TRUE* and *primaryOnly=TRUE*. A Principal Component Analysis was then applied to normalized read counts from *DESeq2 vst* ^54^ method to investigate major axes of variation in the expression data.

Gene-level counts were used for testing of differential expression between *vgll3* genotypes, sampling dates, and their interaction using a Generalized Linear Model approach with normalization factors for unwanted variation as follows. First, genes were filtered for those expressed using *edgeR* ^55^ function *filterByExp* and defining treatment groups. A “naive” GLM was defined with *vgll3* genotype, sampling date (scale-normalized), and their interaction, using *DESeq2*. Non-differentially expressed genes based on Wald’s test and a significance threshold of 0.05 (adjusted for multiple testing) between genotypes, sampling dates and their interaction were extracted using contrasts and the *results* function. The non-differentially expressed genes were designated as “negative control” genes to be used in estimating unwanted variation in the data. For calculating factors that describe unwanted variation, RNA-seq counts were adjusted for differential sequencing depth using the *EDASeq* package ^56^ and the function *betweenLaneNormalization*. RNA-seq read counts for negative control genes were then used to calculate a sample-specific adjustment factors (weights) corresponding to the maturity trajectory of each sample using the *RUVg* method ^34^ and parameter *k=2*. A final GLM was defined with *vgll3* genotype, sampling date (scale-normalized), their interaction, and the two normalization factors from *RUVg*. The model fit for all factors were tested against null models using the *DESeq* function with parameter *test=“LRT”*. Inclusion of all factors significantly improved the fit of the model and were therefore retained in the final analysis. Differentially expressed genes were extracted using Wald’s test and contrasts similar to the naive analysis above.

Gene annotations for gene sets were fetched using NCBI Batch Entrez search and GO term over-representation was tested against expressed genes with *AnnotationHub* ^57^ (snapshot date 2023-04-24) function *enrichGO* and a False Discovery Rate threshold of 0.1.

Gene co-expression networks were constructed using *WGCNA* ^58^. Gene-level counts were normalized using RUV weights and *vst* function of *DESeq2*. Gene co-expression modules were identified using the automatic network construction and module detection function *blockwiseModules* in *WGCNA R* package. A soft-thresholding power of 10 was selected based on best fit of scale-free topology while conserving a moderately large module size. Additional parameters for *blockwiseModules* were defined as *maxBlockSize=10000*, *TOMType = “signed”*, *minModuleSize = 25*, *reassignThreshold = 0*, and *mergeCutHeight = 0.3*. Module neighborhood statistics were analyzed and visualized using the *igraph* R package ^59^. Module genes were tested for overrepresentation of GO terms as described above.

Difference in rank-transformed module eigengene expression between *vgll3* genotype was tested using a linear model (*lm* function) and *p-*values were corrected using the FDR method of ref ^60^. Correlation between module eigengenes and sampling date was calculated using *bicor* function in *WGCNA*. Significance of associations was tested using *corPvalueStudent* function and corrected using the FDR method of ref ^60^.

### ChIPmentation analysis

ChIPmentation reads were quality-trimmed using *fastp* and parameters *--low_complexity_filter --trim_front1=2 --trim_front2=2*. Replicate sequencing runs for the same samples were trimmed individually and trimmed reads were combined into a single *fastq* file. Reads were then aligned to the Atlantic salmon genome downloaded from Ensembl (Salmo_salar-GCA_905237065.2) using *Bowtie2* ^61^ and parameters *--very-sensitive --maxins 1500 --end-to-end*. *Samtools view* was used to filter for primary alignments with mapping quality score over 20 (*-F 256 -q 20*). *Picard MarkDuplicates* ^62^ was used to identify and remove duplicate reads. ChIPmentation fragment coverage was combined for all samples and for replicate samples across genotypes using *samtools merge*. Combined ChIPmentation coverage was downsampled to be equal in genotypes using *Picard DownsampleSam*. We used *MACS3* (https://github.com/macs3-project/MACS) with parameters *--broad --broad-cutoff 0.1* to identify genome regions associated with H3K27ac, H3K4me3 and VGLL3 (for all samples and for *vgll3* genotypes separately). Custom R code and *bedtools intersect* ^63^ were used to filter out peaks overlapping 1 kilo base windows with top 1% of sequencing coverage of control (tagmentation) libraries in order to exclude problematic genome regions. Peaks were tested for over-represented transcription factor binding motifs using HOMER ^64^ and the function *findMotifsGenome.pl*, specifying the parameters *-size given* and *-mset vertebrate*. Peaks were filtered for those with log fold-change compared to input >2 and overlaps of peaks were analyzed in R. Peaks were assigned to closest expressed transcripts in R using the list of expressed genes from RNA-seq analysis. Transcripts were assigned to genes using *biomaRt* function *getBM* ^65^ and annotations were fetched using NCBI Batch Entrez. GO overrepresentation of associated genes was tested as above against background sets defined in the main text.

### *vgll3* knockdown in organ culture

Paired testes from one year old immature males were dissected using fine forceps (FST, Germany) into phosphate-buffered saline (PBS), then stored on ice in Dulbecco’s phosphate-buffered saline (D-PBS, containing 1 mM CaCl_2_ and MgCl_2_) until transferred to culture. Testes were cultured as whole organs at the air-liquid interface on Corning® Transwell® polycarbonate membrane cell culture inserts in polystyrene dishes filled with culture medium. The pore size of the polycarbonate membrane inserts was 0.4um. Basal culture medium consisted of Leibovitz-15 (Invitrogen) supplemented with 10 mM HEPES (pH 7.4), 0.5% w/v bovine serum albumin fraction V (Roche), 0.4 mg/L amphotericine B (Fungizone®; Invitrogen). Retinoic acid was omitted from the culture medium. VGLL3 antibody was added to the culture medium to 6.3 ul/ml and the samples were incubated for 3 days before changing to culture medium without antibody. Control cultures were supplemented with BSA. Samples were cultured for 8 days at 13-14 °C.

#### Fixation and processing

Testis organ cultures were briefly immersed in 100% methanol to prevent coiling of the tissue. For each pair of organ or tissue cultures, one testis was fixed in 4% paraformaldehyde in PBS (4% PFA) and the other testis was fixed in a formol-alcohol fixative (Lillie 1965; ethanol, formaldehyde, glacial acetic acid). Fixation was performed overnight at 4 °C with rocking. Fixed specimens were processed to 100% ethanol for storage at −20 °C. From 100% ethanol, testis cultures were cleared in xylene, followed by a 1:1 xylene:paraffin wax step, and transferred to low-melt paraffin wax, which was changed and incubated overnight at 58 °C. Finally, cultures were embedded in fresh low-melt paraffin wax, sectioned at 7 μm, and attached to SuperFrost® Plus microscope slides. For histological analysis, sections were stained using the periodic acid Schiff (PAS; Sigma-Aldrich) staining reaction and mounted with DPX mounting medium (Sigma-Aldrich). Sections were imaged using an Olympux BX63 microscope.

## Supplementary material

### Supplementary analyses

#### *vgll3* genotype associates with maturation timing in controlled conditions

We tested the association between *vgll3* genotype and maturation probability in controlled settings in 1,5 year-old fish. We used a generalised linear animal model approach to calculate the probability of males reaching maturity in their second year based on their *vgll3* genotypes, similar to ref ^24^. In summary, the animal model was run with a probit-link function under Bayesian Markov Chain Monte Carlo simulations as implemented in the *R* package *MCMCglmm* ^66^. We used a model *y* = *vgll3* + *a + t +e* where *y* is a vector of maturation binaries, *vgll3* is the genotype, *a* is the additive genetic relationship-matrix predicted animal effect ^67^, *t* is the common environmental tank effect and *e* the error term (with fixed variance of 1). Priors were chosen from a distribution following ref ^68^ and based on inferences on 10,000 samples collected every 1,000 iteration after burn-in of 10,000. Model convergence was tested using Heidelberger and Welch’s diagnostic ^69^.

Resembling the earlier maturation age of wild Atlantic salmon carrying the *E* (*early*) allele of *vgll3* ^14^, the *vgll3EE* genotype had a higher maturation probability compared to the *LL* (*late*) genotype, with the *EL* genotype having a probability intermediate between the two (Fig S1). The differences were not statistically significant, likely because of the overall high maturation occurrence during the second year, and smaller sample sizes due to the large size of the fish. Using the same families and experimental set up, assayed during their first year of growth, we had previously showed that the *vgll3EE* genotype has statistically higher maturation probability compared to the *vgll3LL* genotype ^24^. The effect of *vgll3* genotype on maturation probability therefore seems stable across different age-classes.

**Figure S1.**
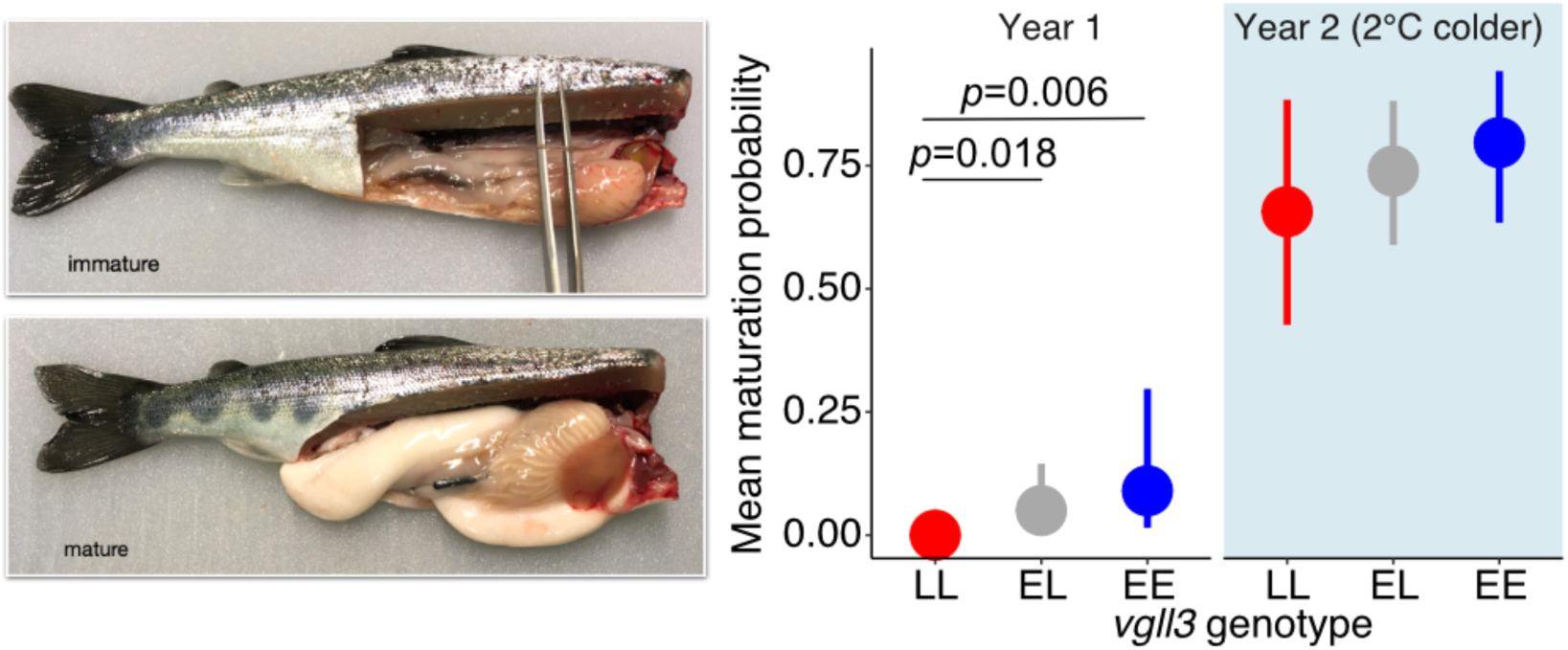
Association between *vgll3* genotype and maturation probability was studied in salmon males raised in controlled conditions. Results for year 1 maturation were previously published in ref ^24^. Fish measured at year 2 were raised at 2 degrees colder temperature from fertilization onwards. *P*-values are shown only for significant contrasts.

### Factor analysis of RNA-seq data

We accounted for uncontrolled variation in testis gene expression by using a factor analysis approach. The reasoning for this approach was that uncertainty regarding the maturation trajectory affects disproportionately those samples that were acquired early in the season; immature testes acquired during breeding season have a higher probability of being truly immature compared to samples acquired earlier. Indeed, examination of major sources of variation in testicular gene expression before factor analysis pointed towards structure in the data largely unrelated to *vgll3* genotype or season. Genes with strongest influence on principal component 1 (top 5% of PC1 loadings) were over-represented in cell cycle, replication and division processes, while those with strongest influence on PC2 were over-represented in cell cycle control and cellular cytoskeleton, among others. The results were overall consistent with known transcriptional changes in maturing salmon testes ^70^. The first principal component correlated strongly with *cyclin B1* expression; immature samples acquired late in the season showed low *cyclin B1* expression, indicating that they were characterized by cell-cycle arrest at the G2/M phase, while some early season samples showed high *cyclin B1* expression consistent with active cell proliferation. The second principal component correlated strongly with the expression of *AMH* (third highest loading on PC2 among all expressed genes, linear model r2=0.53 *p*=1.16e-04), a marker for maturation trajectory in salmon testes ^24^, with some early season samples showing lower expression. This is expected if these samples were progressing towards initiating maturation. In other words, the results indicate that large-scale variation in the data could be explained by sample distribution across a maturation continuum, where, despite being morphologically immature, samples varied in their molecular state with respect to cell cycle and maturation initiation.

We performed factors analysis ^34^ to account for this unwanted variation in maturation stage. We calculated normalization factors based on genes not differentially expressed in a first-pass general linear model (GLM) analysis with *vgll3* genotype and sampling date as effects. These normalization factors (K=2) were strongly correlated with an alternative normalization approach where we identified unsupervised surrogate variables ^71^ orthogonal to the effects of *vgll3* genotype and sampling date (Pearson *r*=0.95). The normalization factors were then included in a final GLM model along with *vgll3* genotype, sampling date, and their interaction. The final GLM model performed significantly better to identify a broader set of DEGs that largely included the DEGs identified using the naive model without normalization factors. These results indicate that our approach robustly accounted for variation in the maturation stage of the samples and increased the power to identify relevant gene expression differences.

### VGLL3 association with Hippo-Ras genes

We investigated the association between gene co-expression modules and VGLL3 binding to identify signaling pathways putatively controlled by VGLL3. We asked whether *tead3* (Hippo) network neighbors in the *magenta* module were associated with *vgll3* binding, as expected if *tead3* co-expression with these genes was mediated by VGLL3. Of the 24 *tead3* network neighbors, 14 had VGLL3 binding regions in close genomic proximity (4 with VGLL3 promoters, 6 with VGLL3 enhancers), including the negative regulator of Ras signaling, *neurofibromin* (*NF*, Fig S2). Two additional Ras GAPs (GTPase activating proteins) not included in *magenta* showed proximal VGLL3 peaks (*RASA1*, *RASAL1,* Fig S2). *Tead* has been show to directly determine Ras signaling activity in *Drosophila* through regulation of the Ras default inhibitor *cic* and activator *ets1* (a.k.a. *Pnt*), thus controlling for the balance of cell proliferation versus differentiation ^72^. We therefore investigated VGLL3 binding on the seven *cic* and *ets1* gene copies (not included in *magenta,* one *cica* gene included in *blue*) in Atlantic salmon and found all to be associated with strong VGLL3 peaks. Together, the results strongly suggest that Hippo-VGLL3 mediated control of cell proliferation in the testes is mediated in part by their effect on overall Ras/MAPK signaling activity, as well as its effectors. *Vgll3*-mediated regulation of Ras genes provide a putative link for mediating *vgll3*-associated oncogenic development as aberrant Ras signaling is one of the most frequent causes cancer ^73,74^.

**Figure S2.**
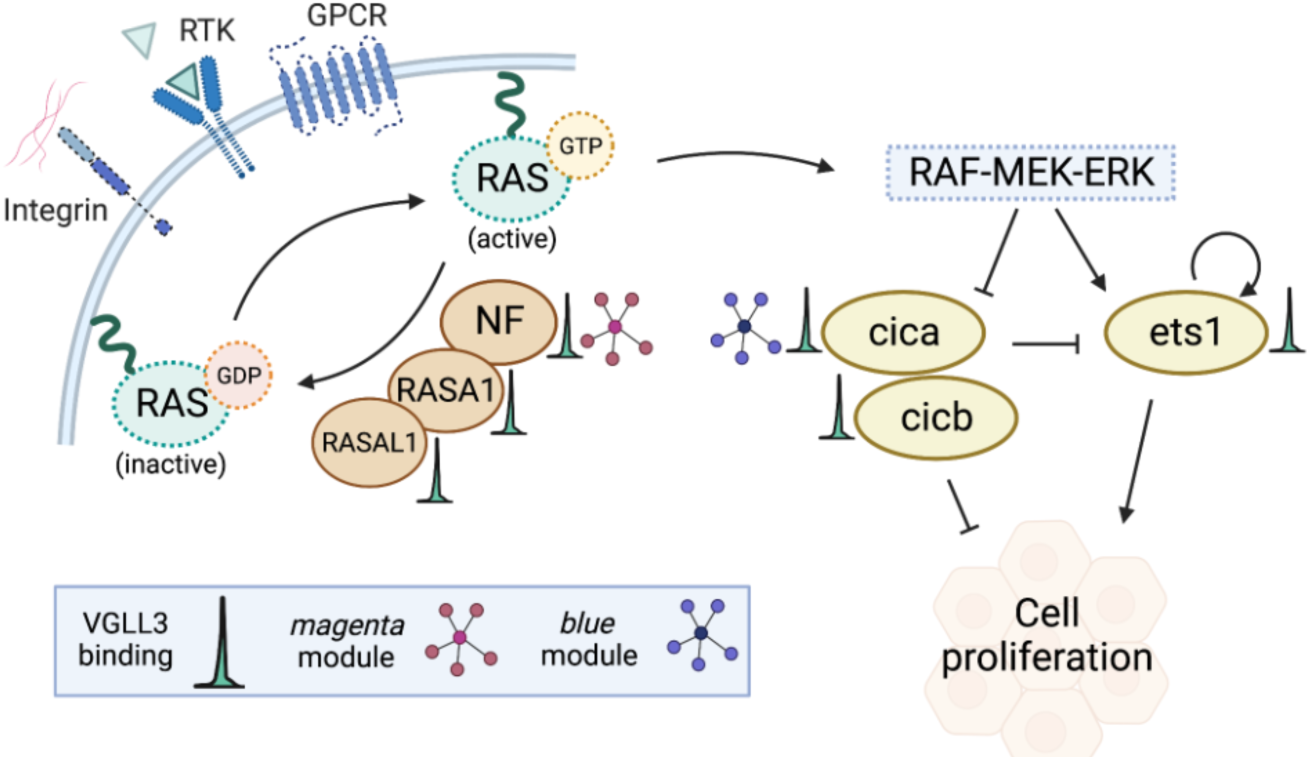
Ras signalling pathway genes show functional association with *vgll3*. Ras/MAPK pathway is molecularly connected to *vgll3* through co-expression modules (magenta and blue) as well as VGLL3 binding regions in proximity of Ras inhibitors *NF*, *RASA1*, *RASAL1*, as well as Ras effectors *cica*, *cicb*, and *ets1*. Ras responds to signalling input from the extracellular matrix (integrins), growth factors (RTK, receptor tyrosinase kinase), and G-protein coupled receptors (GPCR) to control cell proliferation versus differentiation. Genes are annotated with VGLL3 binding peaks and inclusion in co-expression modules.

### Supplementary figures

**Figure S3.**
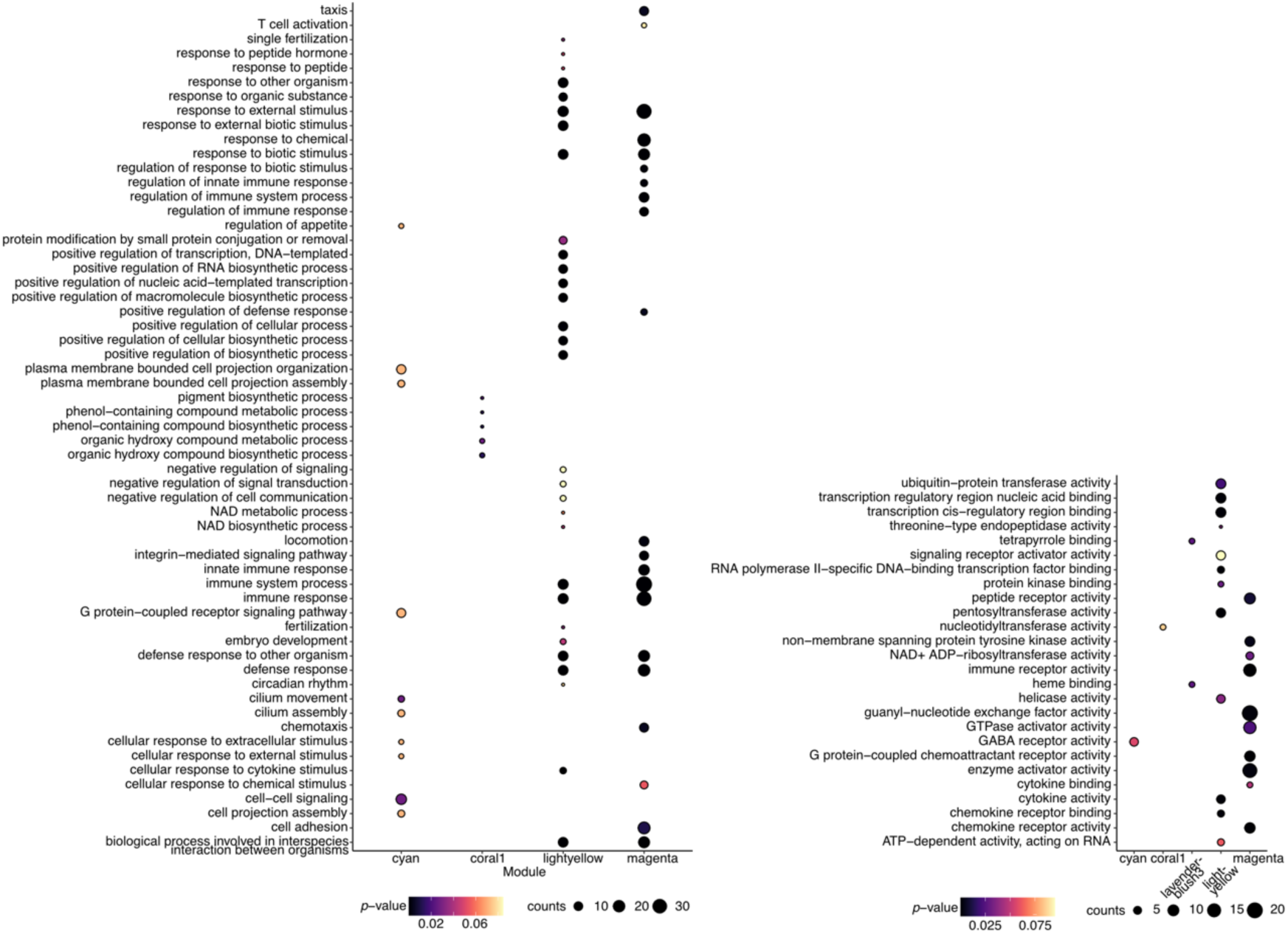
Gene Ontology (left panel: biological process, right panel: molecular function) over-representation for modules associated with *vgll3* genotype.

**Figure S4.**
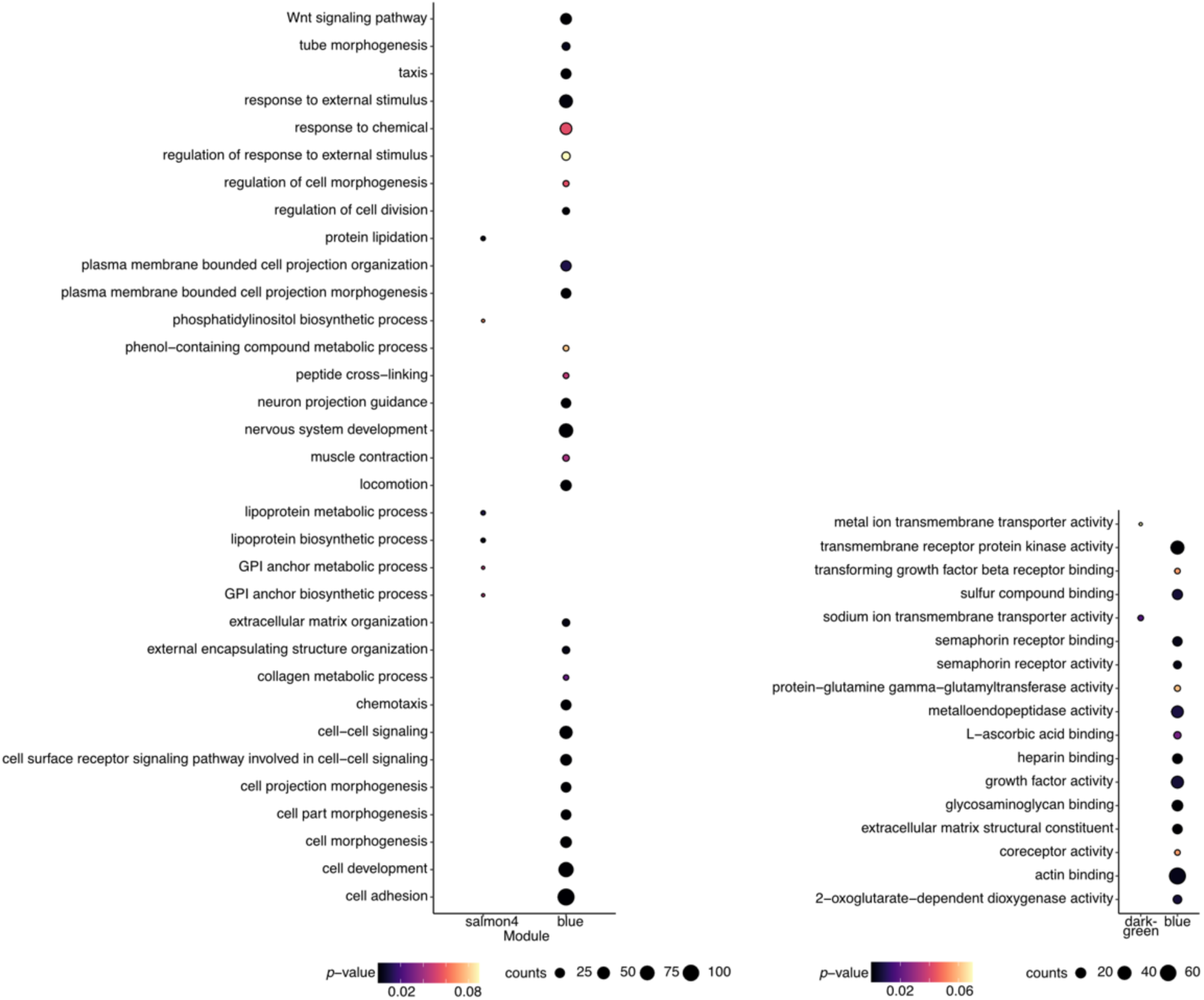
Gene Ontology (left panel: biological process, right panel: molecular function) over-representation for modules associated with sampling date.

**Figure S5.**
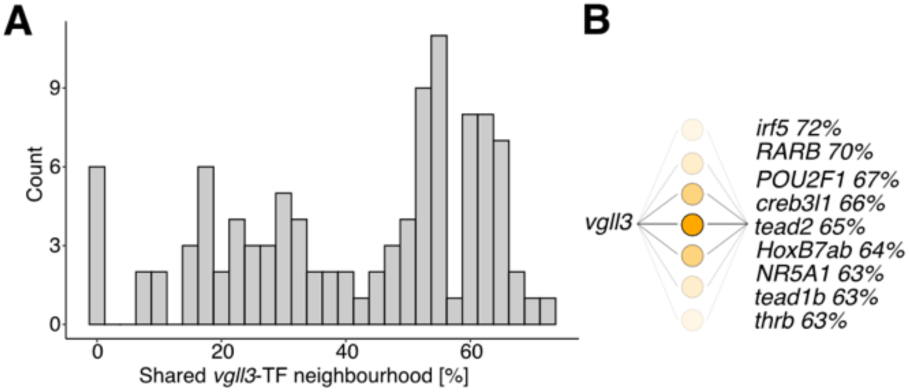
Network neighborhood sharing between *vgll3* and transcription factors in *blue* module. A) The percentage of share neighbors between *vgll3* and 106 transcription factors included in *blue*. B) Transcription factors sharing most blue neighborhood represent diverse signalling pathways and include, e.g., *tead2* and *nr5a1*. Transcription factors are ordered according to the percentage of shared network neighborhood.

**Figure S6.**
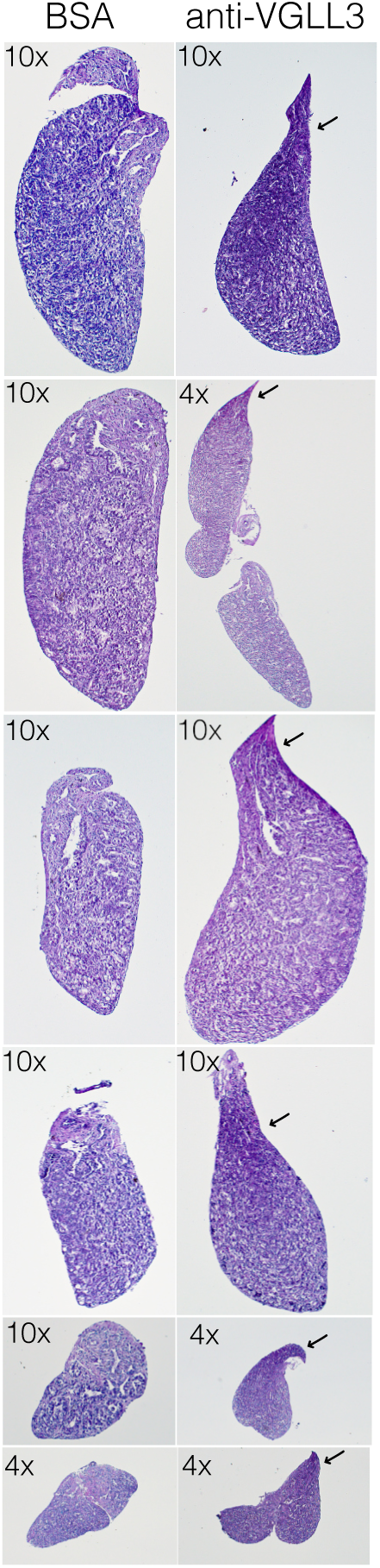
VGLL3 knock-down in ex-vivo organ culture. Paired testes from single individuals are cultured in control conditions (BSA) or in the presence of anti-VGLL3 antibody. Sections show increased cell proliferation and motility in VGLL3 knock-down conditions.

**Figure S7.**
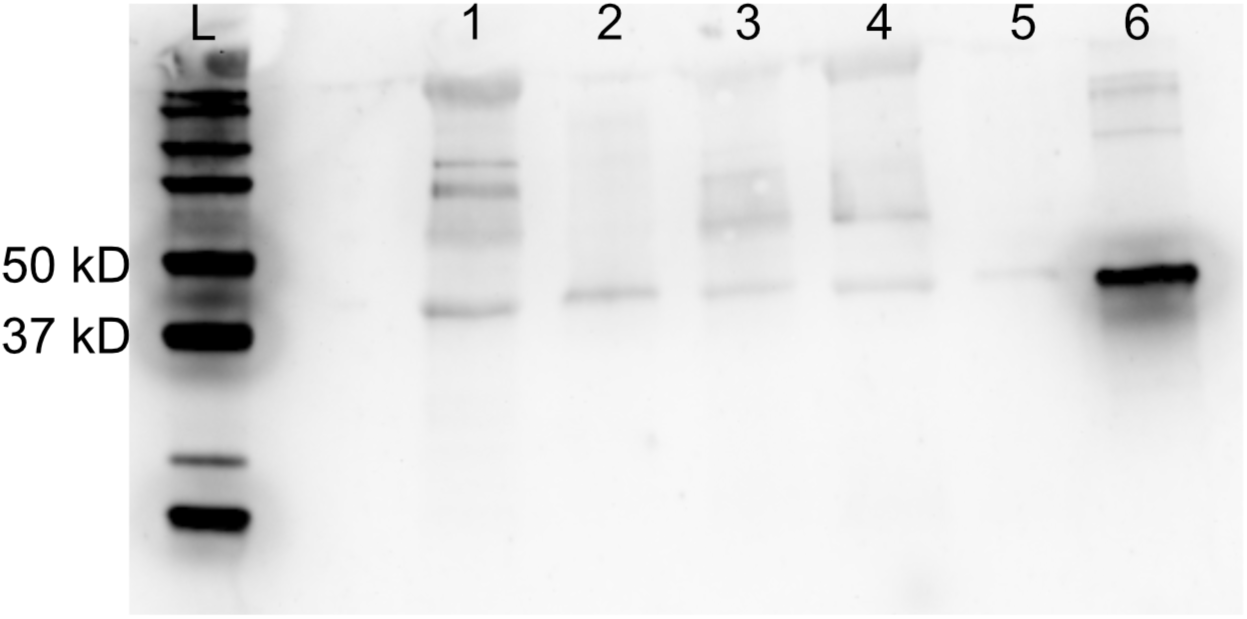
Validation of anti-VGLL3 antibody. Co-immunoprecipitated samples and controls were assayed using Western blot and anti-VGLL3 antibody. Estimated molecular weight of VGLL3 protein is 42 kD. L, ladder; 1, sample 1 (heart tissue); 2, 0.05 ug of VGLL3 protein; 3, sample 2 (heart tissue); 4, sample 3 (heart tissue); 5, 0.01 ug VGLL3 protein; 6, 0.005 ug of VGLL3 protein (without co-immunoprecipitation).

### Supplementary tables

Table S1. DEGs with significant effect of *vgll3* genotype, sampling date (season), and their interaction.

Table S2. Alignment statistics for multiomic sequencing data.

## Notes

### Competing Interest Statement

The authors have declared no competing interest.

